# BitBIRCH-Lean: chemical space in the palm of your workstation

**DOI:** 10.1101/2025.10.22.684015

**Authors:** Ignacio Pickering, Krisztina Zsigmond, Kenneth López Pérez, Miroslav Lžičař, Ramón Alain Miranda-Quintana

**Affiliations:** Department of Chemistry & Quantum Theory Project, University of Florida, Gainesville, Florida 32611, USA; Deep MedChem a.s., Revoluční 764/17, Prague 1 - Old Town, Czechia, IČO: 21029962

## Abstract

We present BitBIRCH-Lean, a fast, memory-efficient implementation of the BitBIRCH algorithm, designed for high-throughput clustering of huge molecular libraries (up to billions of drug-like molecules) on typical workstations. BitBIRCH-Lean considerably improves on the original BitBIRCH implementation by incorporating dynamic types and bit-packed fingerprints inside the clustering tree. Most operations in BitBIRCH-Lean are efficiently performed on compressed data, and optional C++ extension accelerate the bottleneck calculations, providing up to 2X speedup. Benchmark tests against GPU-accelerated methods highlight BitBIRCH-Lean as an efficient alternative for processing vast amounts of molecules. We further demonstrate the versatility of this new package by showcasing a parallel, multi-round variant of the BitBIRCH algorithm that exploits the gains in efficiency to cluster hundreds of millions of molecules in minutes, with no loss in cluster quality. The code is freely available at: https://github.com/mqcomplab/bblean.

## 1 Introduction

Clustering has long been a cornerstone of cheminformatics, ^1,2^ drug design,^3,4^ and molecular modeling,^5,6^ enabling the systematic organization of molecular information into chemically meaningful subgroups. By partitioning molecules according to structural similarity, clustering facilitates the identification^7,8^ of scaffold families, guides compound library design, and highlights trends relevant to biological activity. In the context of drug discovery, clustering supports lead identification by reducing chemical redundancy and ensuring that screening^9,10^ campaigns sample diverse chemotypes rather than over-representing closely related analogs. Cluster centroids can then be used in virtual screening, molecular docking, or pharmacophore modeling, thereby reducing computational costs while retaining coverage of the underlying chemical space. Beyond these direct applications, clustering aids in the visualization of complex datasets^11,12^ (even allowing linear projection methods to surpass non-linear alternatives), helps interpret high-dimensional molecular fingerprints, and serves as a foundation for machine learning workflows that depend on representative or balanced training sets. Collectively, these roles and applications underscore the centrality of clustering to modern computational drug design and cheminformatics.

Despite these key roles, the utility of clustering has been constrained by the limitations of conventional algorithms when applied to modern chemical libraries. Public databases such as PubChem^13^ and ChEMBL^14,15^ already contain millions of molecules, while commercial and virtual collections such as Enamine REAL^16^ now enumerate tens of billions of synthetically accessible compounds. In addition, *de novo* molecular generation^17,18^ methods routinely propose virtual chemical spaces that reach trillions of structures. Standard clustering approaches, including hierarchical agglomerative clustering, ^19–21^ the Butina^22–24^ algorithm, and density-based approaches such as DBSCAN,^25–27^ have proven useful at modest scales but exhibit unfavorable scaling with respect to time and memory. Hierarchical methods offer chemically interpretable trees but suffer from quadratic or worse complexity. Butina clustering, perhaps the most widely used method in cheminformatics, avoids explicit hierarchy construction but still requires repeated all-against-all similarity calculations, limiting its use to datasets of at most a few million molecules. Partition-based methods such as k-means^28^ scale better in principle but are poorly suited to sparse binary fingerprints such as ECFP,^29^ which dominate modern cheminformatics, and tend to impose spherical cluster assumptions that are chemically unrealistic.

To address these challenges, we recently proposed BitBIRCH,^30^ a novel clustering algorithm tailored to the unique properties of binary molecular fingerprints and designed explicitly for large-scale applications. BitBIRCH circumvents the quadratic scaling barrier by combining the instant similarity (iSIM) ^31–33^ framework together with a tree structure to efficiently store and traverse the fingerprints. Remarkably, BitBIRCH achieves linear-time behavior relative to the number of molecules, while preserving the chemical interpretability and resolution of clusters. BitBIRCH delivers robust clustering results that maintain fine-grained distinctions between scaffolds and analog series, making it attractive as a tool for scaffold hopping, diversity analysis, and virtual screening. However, the memory requirements of the original implementation made it difficult to fully exploit the algorithmic advantages of BitBIRCH.

In this contribution, we introduce BitBIRCH-Lean, a package based on the original BitBIRCH implementation, but greatly surpassing its efficiency. By completely restructuring how fingerprints are handled, and optimizing how individual molecules are passed through the tree, we achieve remarkable improvements. Dynamically assigning the data type required to safely store each subcluster means that we can start processing much more amenable *uint8* arrays. Additionally, packing the fingerprints not only saves more memory, but also improves time efficiency. These improvements make it possible to process hundreds of millions of molecules with minimal hardware requirements. We leverage this by discussing a simple recipe to simultaneously process multiple batches of fingerprints in a novel pipeline that allows clustering hundreds of millions of molecules in a single workstation. The code and examples associated with BitBIRCH-Lean can be found in https://github.com/mqcomplab/bblean.git.

## 2 Methods

### 2.1 Overview of the BitBIRCH algorithm and effective refinement strategies

We present here a summary of the main concepts behind the BitBIRCH algorithm, ^30^ and the most effective cluster refinement strategies to obtain high quality clusters. ^34^ For an indepth analysis and explanation of the BitBIRCH procedure we refer readers to the original articles.^30,34,35^

BitBIRCH is based on two key ideas: using a tree structure to help storing and traversing the clustered fingerprints, and leveraging the iSIM framework to conveniently calculate cluster properties. The tree is composed of two distinct structures: nodes and subclusters, with nodes functioning as containers for the subclusters. A user-defined branching factor caps the maximum number of subclusters that a node can store. This fixed maximum capacity per node is critical to avoiding a *O*(*N* ^2^) exploration. When an incoming molecule cannot be accommodated in a node’s current subclusters, it acts as the starting point for a new subcluster unless the maximum node capacity has been reached, which triggers a node split. Node splits create two new nodes from a starting node around maximally-separated subclusters, causing the tree to increase in depth by one level.

Subclusters store information in bit features (BFs), which contain: the number of molecules in the cluster (**N**), a vector with indices of the clustered molecules (**mol**), and a vector which holds in its *k*^*th*^ component the sums of the bits in the k^th^ position for all fingerprints in the cluster (**ls**). In principle, these are the minimal structures required to specify a given subcluster, but subclusters additionally store the cluster centroids (**c**) in order to avoid on-the-fly recalculation, which is required when evaluating merge candidates.

**N** and **ls** provide all data needed to calculate the iSIM of the cluster, which corresponds to its diameter-complement. BitBIRCH accommodates several criteria to decide when to merge two subclusters: by the diameter or radius of the resulting cluster (the latter is also easily accessible through iSIM-like manipulations). It is also possible to have tighter merging criteria, by using the tolerance option, which enforces the quality (radius or diameter) of an already-formed cluster to not decrease by a pre-specified user parameter. The serial tests discussed below use the diameter merge option, and in the parallel pipeline we also include tolerance in the merge to ensure the overall quality of the final partition.

As with other “radial-based” clustering methods, a single BitBIRCH pass sometimes generates a large, overly-populated cluster. We recently proposed multiple ways to refine these results and avoid this issue. In the spirit of keeping this new implementation as simple as possible, we only retained one of these strategies: BF refinement. The corresponding workflow is as follows: (a) Cluster all the molecules and identify the largest cluster. (b) Split the largest cluster and separate all molecules into single-element BFs (all remaining BFs are kept as they are). (c) Re-cluster all BFs collected on the previous step, optionally using a stricter merge criterion to maintain the distribution of average Tanimoto similarities.

### 2.2 Implementation and software library organization

BitBIRCH-Lean contains a re-implementation of all functionality in the original BitBIRCH codebase, with added usability, memory-efficiency, and performance improvements. The original BitBIRCH software loosely followed the *scikit-learn* ^36,37^ application programming interface (API); we chose to maintain this interface for BitBIRCH-Lean, expanding it to fully implement all *scikit-learn* functions, so the software could be integrated with existing workflows with minimal modifications.

Besides its Python API, BitBIRCH-Lean includes a simple command line interface (CLI) that exposes its parallel and serial implementations as direct commands, together with fingerprint manipulation utilities (splitting and merging fingerprint files and creating packed bit-packed fingerprints starting from SMILES strings), tools for generating useful visualizations^38–40^ and basic cluster analyses. The CLI is meant for quick exploration of molecular libraries; it requires no code to perform a full clustering pipeline starting from a SMILES file.

## 3 Results and discussion

For all clustering workflows, datasets consist of tranches randomly chosen from the ZINC22^41^ database. All benchmark runs were performed on a workstation with an Intel Core i9-10980XE 3.00 GHz CPU with 18 physical cores and Hyper-Threading enabled (36 logical cores), running Ubuntu 24.04 (Linux 6.8, GLIBC 2.39), using Numpy 2.3.2^42^ and Python 3.11.13. Memory usage is reported as the total resident set size (RSS) of a BitBIRCH-Lean process instance together with all its child processes, and was monitored with a 10 ms interval. The C++ BitBIRCH-Lean extensions were compiled with GCC 12.4, and use the *-O2, -march=nocona, -mtune=haswell* optimization flags.

All GPU-accelerated *nvMolKit* ^43^ calculations use an NVIDIA GeForce RTX 5090 GPU (Blackwell architecture, compute capability 12.0) with 32 GiB VRAM. *nvMolKit* (beta version 0.1) was compiled against the CUDA Toolkit version 12.9.

The BitBIRCH-Lean code is available at: https://github.com/mqcomplab/bblean.

### 3.1 Serial, Python-based BitBIRCH-Lean

In order to make the use of iSIM in the original BitBIRCH as simple as possible, the fingerprints that went through the tree had to be converted to what we term *dense* (one integer per feature) *int64* (64-bit signed integer) arrays. This is due to the **ls** vector, which needs to store the sum of an *a priori* unspecified number of fingerprints for each cluster. This design choice greatly simplified the original implementation of BitBIRCH, but placed a considerable burden in the memory required to run the method (e.g. storing one million fingerprints, each with 2048 bits, as *int64* demands 16 GiB RAM). BitBIRCH-Lean bypasses this issue with two key modifications: dynamic integer types for all arrays stored in the tree, and using bit-packed instead of dense arrays whenever possible. In what follows we explain these two methods in more depth.

Dynamic type assignment relies on overflow checks performed each time fingerprints are added to the tree, and casting the corresponding arrays to the minimum unsigned integer dtype that can safely hold the result of the sum. The checks represent a negligible overhead, and the casts are triggered only sparingly, when a cluster grows to a large size.

Bit-packing *uint8* fingerprints requires mapping single bytes to single bits within a sequence of bytes. In practice, native Numpy^42^ operators are used to bit-pack fingerprints produced or loaded as *uint8*, reducing the per-molecule footprint from *F* bytes to ceil(*F/*8) bytes (e.g., a 2048-bit ECFP occupies 256 bytes), with any necessary zero padding handled automatically.

All hot paths (intersection/union counts, iSIM operations, centroid maintenance, and node splitting) operate directly on these compact bit-packed buffers via Numpy bitwise operations. Additionally, when the fingerprint length is a multiple of 64, the buffers are reinterpreted as *uint64* to exploit 64-bit hardware *popcount* intrinsics.

Subcluster centroids remain packed as well: we maintain a running vector of per-bit occurrence counts, and the majority-vote centroid is repacked before persistence. Compared with dense (byte-per-feature) layouts, operating on packed buffers reduces memory traffic, keeps more of the tree’s working set in cache, and decreases computational cost. Beyond these critical changes, the BitBIRCH-Lean codebase contains multiple other optimizations to further increase the clustering performance.

In terms of performance increase, the new Python BitBIRCH-Lean implementation is on average 2.20x faster than the original implementation. Performance benchmarks are shown in Figure 2, which demonstrates that the performance gains are scalable over a range of 1M fingerprints. Bit-packing alone yields an eightfold reduction in memory, and the additional dynamic typing and extra optimizations results in an overall reduction of 30x in RAM usage, as shown in Figure 1, where benchmarks were performed, again, up to 1M fingerprints.

**Figure 1.**
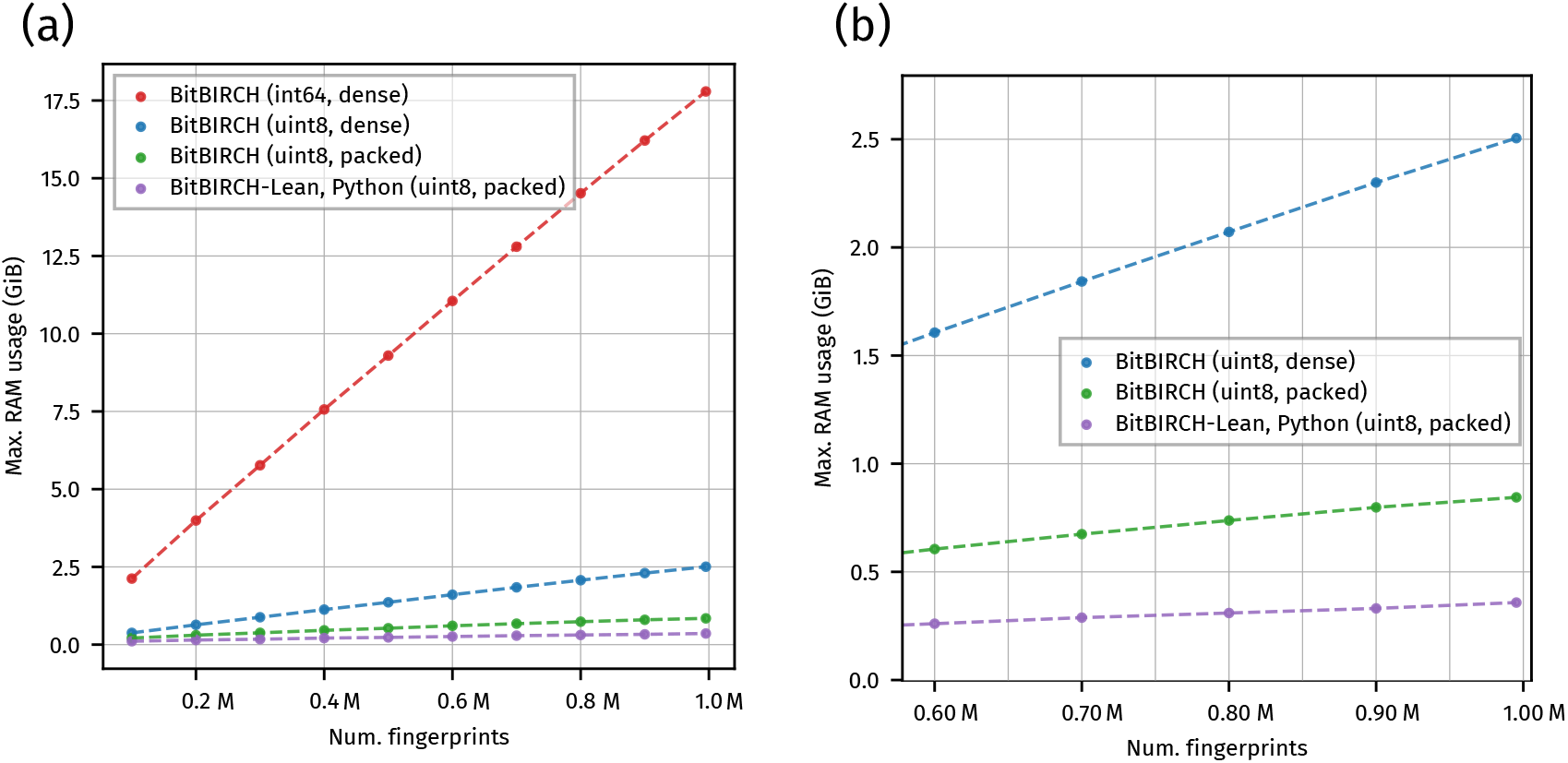
Comparison of RAM usage for different BitBIRCH implementations (branching factor 254), for clustering up to 1M molecules. Traces correspond to the average over three independent runs. Traces labeled *int64* correspond to the original BitBIRCH implementation, while traces labeled *uint8* correspond to a naive use of *uint8* arrays without further code modifications. Dense traces use byte-level features in fingerprints, while packed traces correspond to bit-packed fingerprints. All calculations use a threshold of 0.5. (a) shows a comparison over the full range, while (b) displays a zoomed-in inset focusing on the *uint8* and *Lean* implementations, for the largest sets only.

**Figure 2.**
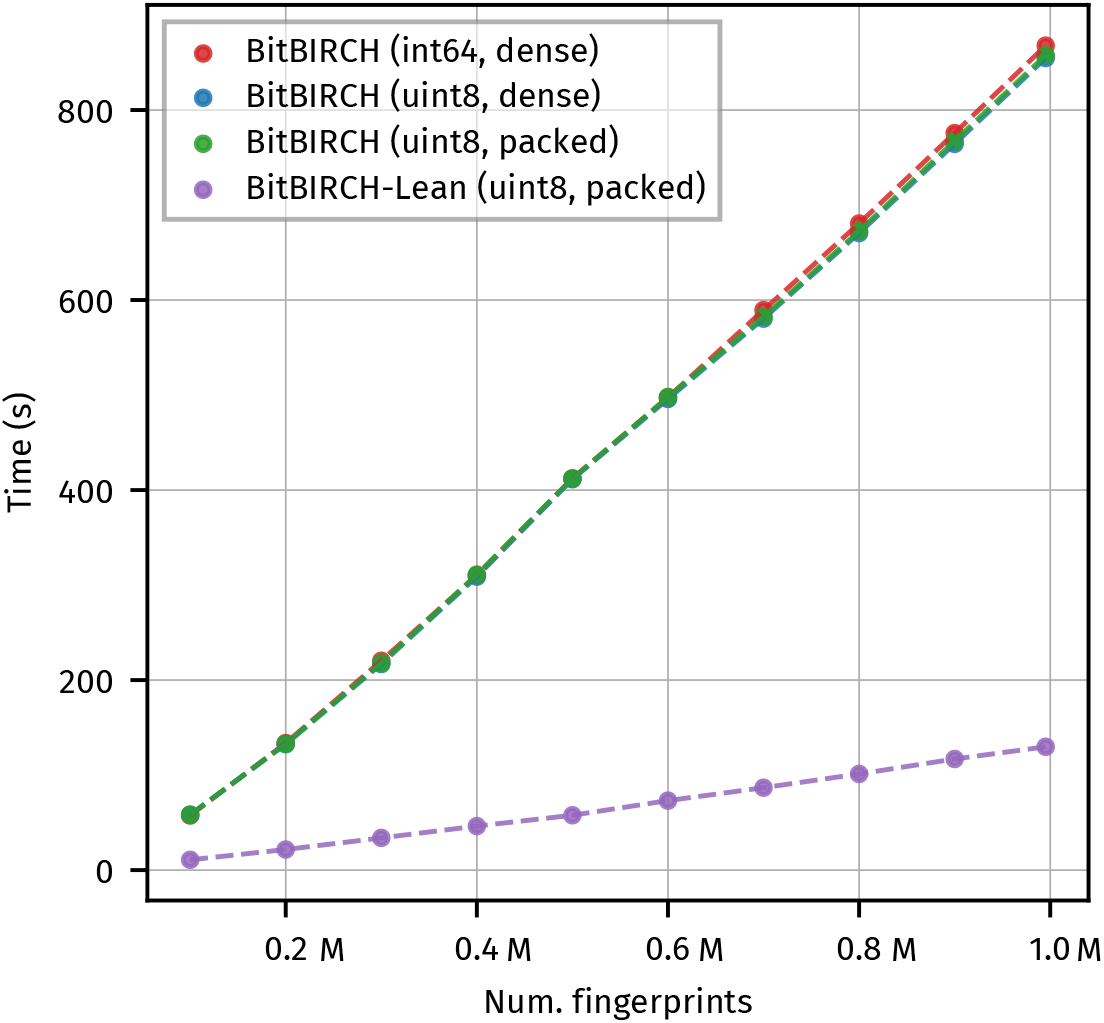
Timings for different BitBIRCH implementations, for clustering up to 1M molecules. Note that there is no significant performance difference when using *uint8* or packed *uint8* fingerprints in the original BitBIRCH implementation, with the corresponding traces almost fully overlapping. Traces correspond to the average over three independent runs. All calculations use a branching factor of 254 and a threshold of 0.5.

For all benchmarks calculations we use a branching factor of 254 and a threshold of 0.5, Analogous plots for a branching factor of 50 can be found in Figures S1^†^ and S2^†^ in the ESI, which show that the performance is not strongly dependent on branching in this range, with larger branching factors resulting in higher gains in RAM usage at a slight performance penalty (still outperforming the original BitBIRCH by approximately the same factors). For specific values reported on these benchmarks see Tables S1^†^ and S2^†^ in the ESI.

### 3.2 Performance of C++ extensions

After careful benchmarking of our BitBIRCH-Lean implementation we identified two remaining bottlenecks in terms of compute performance. The most costly operation was the computation of Tanimoto similarities between incoming fingerprints and the centroids of the BFs held by the tree nodes. The second most costly operation was the calculation of maximally dissimilar BFs within a given node, required for node splitting when the branching factor is exceeded. We implemented both these operations in C++ and dispatched to them through the pybind11^44^ library. This also required a custom implementation of packing and unpacking the bit-packed fingerprints as numpy arrays, and other auxiliary functions such as the iSIM average Tanimoto for a set of fingerprints. Running BitBIRCH-Lean with these C++ extensions resulted in a speedup factor of 2.3x on average compared with Python and Numpy, which is illustrated in Figures S2a^†^ and S2b^†^ in the ESI. This speedup gain is achieved at no additional cost in RAM, as shown in Figure S2c^†^ in the ESI. We note that the overhead of *pybind11* dispatching is negligible compared with the cost of these operations.

Acceleration of the fingerprint-centroids similarity calculations relies on 64-bit popcount intrinsics, so it can be efficiently implemented for fingerprint sizes that are a multiple of 64 only. This is hardly a limitation for fingerprint clustering, since all common chemical fingerprint sizes (512, 1024, 2048, 4096 for ECFPs^29^ and the default *rdkit* fingerprints) fulfill this criterion.

Although we identified opportunities for parallelization using SIMD intrinsics, we currently decided against an explicit SIMD implementation for better maintainability. Only the iSIM average Tanimoto similarities are implicitly vectorized by the GCC and CLang compilers. Further work in this regard is planned for future BitBIRCH-Lean releases.

We believe that for high enough branching factors it may prove beneficial to perform fingerprint-centroid calculations on GPUs. However, the overhead of buffer transfer to an accelerator may require the full algorithm to be implemented on GPU. Detailed benchmarking is necessary to assess whether this would result in performance gains, and this was not explored in this work.

### 3.3 Comparison of serial BitBIRCH-Lean with GPU-accelerated libraries

Recently, interest in implementing GPU-accelerated cheminformatics workflows has increased significantly, partly due to the performance of current GPUs and lower barrier of entry for GPU programming provided by Python bindings to CUDA libraries such as CuPy^45^ and PyTorch.^46^

Since cheminformatics heavily relies on graph descriptors, GPU parallelization of many algorithms can be challenging. The Taylor-Butina^22,23^ algorithm, however, is a prime candidate for this kind of parallelization, since it requires calculating all pairwise Tanimoto similarities within a set of fingerprints. The second stage of the algorithm, the cluster assignment, is less easily parallelizable due to each label assignment step having a sequential dependence on the previous assignments.

The implementation of the cross-Tanimoto Taylor-Butina calculations in *nvMolKit* ^43^ is a recent, highly performant example of this approach. Here we perform benchmarks of the (CPU only) BitBIRCH-Lean Python and C++ implementations, and compare them with a Taylor-Butina clustering pipeline using the CUDA-accelerated *nvMolKit*, followed by a CPU-based label assignment step relying on *rdkit* ^47^ (we name this approach TB-GPU).

Figure 3 compares both workflows in terms of wall time for clustering increasing numbers of fingerprints. For smaller sets, TB-GPU is comparable to BitBIRCH-Lean, but the performance gap with BitBIRCH-Lean widens dramatically owing to the *O*(*N*) scaling of BitBIRCH, and for 62500 fingerprints BitBIRCH-Lean is almost 10x faster. These results show that, even when employing cutting-edge hardware and parallelization, the scaling prefactor for BitBIRCH-Lean is small enough that it outperforms the *O*(*N* ^2^) Taylor-Butina method.

**Figure 3.**
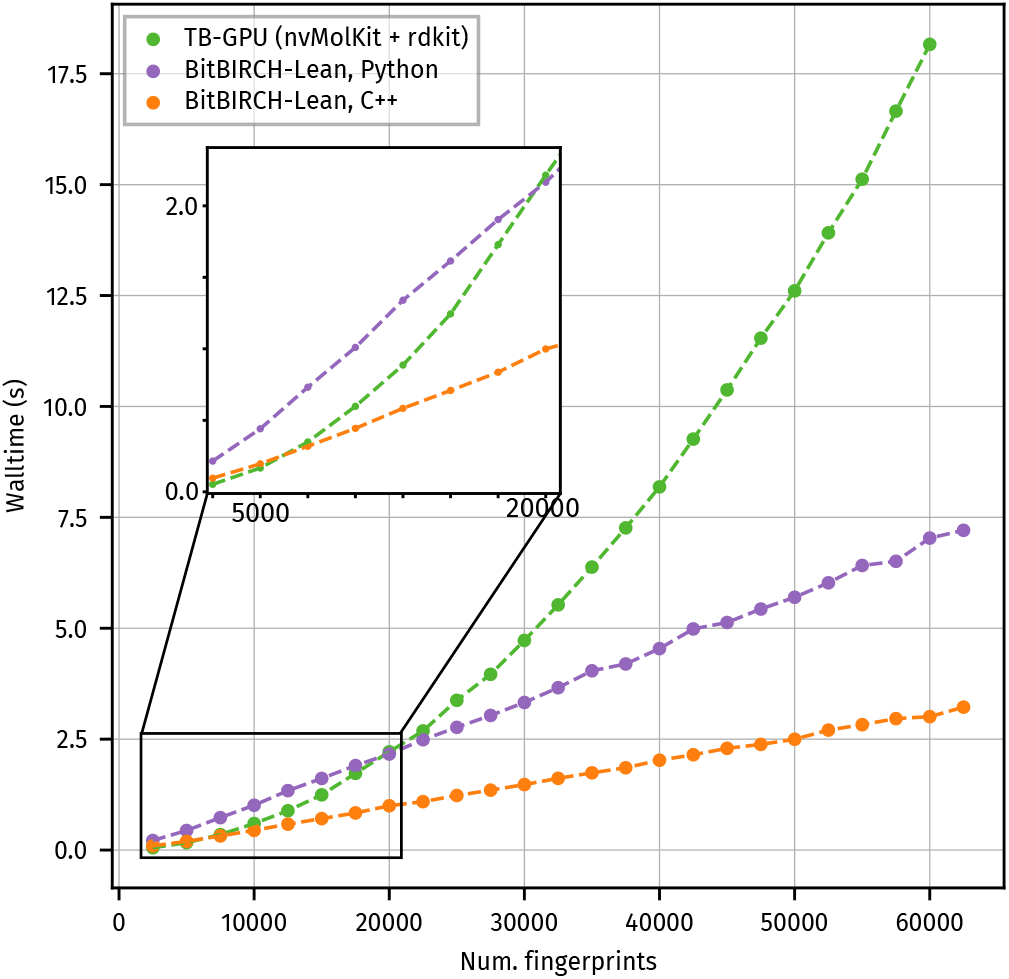
Comparison between the GPU-accelerated Taylor-Butina (TB-GPU) pipeline and BitBIRCH-Lean (Python-only and Python-C++ versions). Timings for sets of fingerprint of sizes ranging between 2500-62500 are shown.

We emphasize that this comparison does not show shortcomings of *nvMolKit* itself, but rather an intrinsic limitation of the Taylor-Butina algorithm. Our comparison is done with respect to the full CPU+GPU clustering pipeline; a comparison of the full BitBIRCH-Lean clustering procedure with the limited GPU-accelerated components of *nvMolKit* is presented in Figure S3a in the ESI^†^, which shows that BitBIRCH-Lean still outperforms the cross-Tanimoto similarity calculation under these conditions, for sets with 15k fingerprints using the C++ extensions and 33k fingerprints using the Python-only implementation. Finally, Figure S3b in the ESI^†^ displays the memory consumption of BitBIRCH-Lean compared with *nvMolKit* (including *rdkit* routines). Memory usage also increases quadratically for both VRAM and CPU RAM, and quickly saturates the available system memory, reaching more than 32 GiB VRAM for 62k molecules. In contrast, BitBIRCH-Lean memory usage scales linearly and remains low throughout (less than 300 MiB).

### 3.4 Parallel, multi-round BitBIRCH-Lean

The standard BitBIRCH algorithm is inherently sequential, since each new fingerprint insertion in the tree influences the cluster in which subsequent fingerprints will be inserted. This fact makes parallelization of BitBIRCH challenging. In the original BitBIRCH implementation,^30^ parallelization was achieved by independently clustering multiple fingerprint files using the *radius* merge criterion, and serial clustering of the centroids of all resulting clusters afterwards. This achieved clusters of medium quality, and had very high peak memory usage.

Our approach in BitBIRCH-Lean is an approximate divide-and-conquer strategy that relies on our findings that the quality of the final clustering results can be improved by *refinement* and *reclustering* operations. ^34^ In this implementation we start with a sequence of packed fingerprint files and cluster them independently, in parallel, using the *diameter* merge criterion by default. We name the result of each independent clustering step a *partial tree* (PT). After this clustering ends, parallel BitBIRCH refines each PT a single time by breaking apart the largest cluster into single elements. The subsequent rounds re-insert these elements when building a new tree from the resulting BFs. This refinement step is also independent, thus requires no synchronization, and increases the quality of the PTs.

In our implementation we rely on worker pools for these parallel steps, where the number of worker processes per step can be tuned depending on the memory (fewer processes working in parallel result in less RAM pressure with longer compute times as a tradeoff).

With the refined PTs built, parallel BitBIRCH partitions the resulting sequence into subsequences of PTs and performs a succession of tree-merging rounds. In each of these, new collated PTs are built starting from all PTs in a given sub-sequence. After a predetermined number *M* of tree-merging rounds, a final merging round collects all PTs into a single tree. The leaves of this last tree are the final resulting clusters.

Tree-merging rounds use a variation on the *tolerance* merging criterion, ^34^ which we term *tolerance-diameter*. Two clusters are merged only if the condition

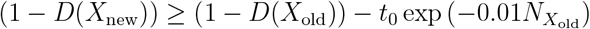

is met, where *t*_0_ is the tolerance value (a free parameter we set to 0.05 by default), *D*(*X*) is diameter of the cluster given by the set of fingerprints *X, X*_new_ is the cluster that would result from the merge, and *X*_old_ is the cluster currently in the tree. This expression guarantees that cluster merging only happens if the resulting cluster is more compact than the inserted one, up to some tolerance that decreases exponentially with cluster size (heuristically, the diameter is an estimate that improves as the cluster size increases).

Diameters use the Tanimoto iSIM approximation, 1 *−* iSIM_JT_(*X*) = *D*(*X*), which for a set with *G* fingerprints, each with *Q* features indexed by *q*, where the componentwise sum of the fingerprints is **k**, is given by:

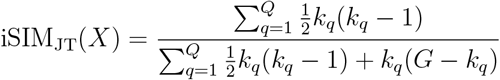

We empirically found this slight variation over the plain *tolerance* criterion to perform better in terms of maintaining the cluster compactness distribution from the initial rounds, and reducing the number of singletons. No comprehensive hyperparameter search was performed to optimize this expression. The same threshold and branching factor as the initial clustering were used in all cases.

The maximum number of PTs in a sequence, *P*, and the number of merging rounds *M*, are additional user-specified parameters that can be tuned to maximize the quality of the final clusters, while taking hardware constraints into account. We provide benchmark results for *P* = 10 and *M* = 1 only, which we found to be relatively robust defaults (except for 1 billion molecules, where *M* = 5 was used). Optimization of these parameters is bound to be system-specific, and depends on the molecular library under consideration.

Figure 4 depicts a schematic diagram of the parallel, multi-round BitBIRCH algorithm, as implemented in the BitBIRCH-Lean software.

**Figure 4.**
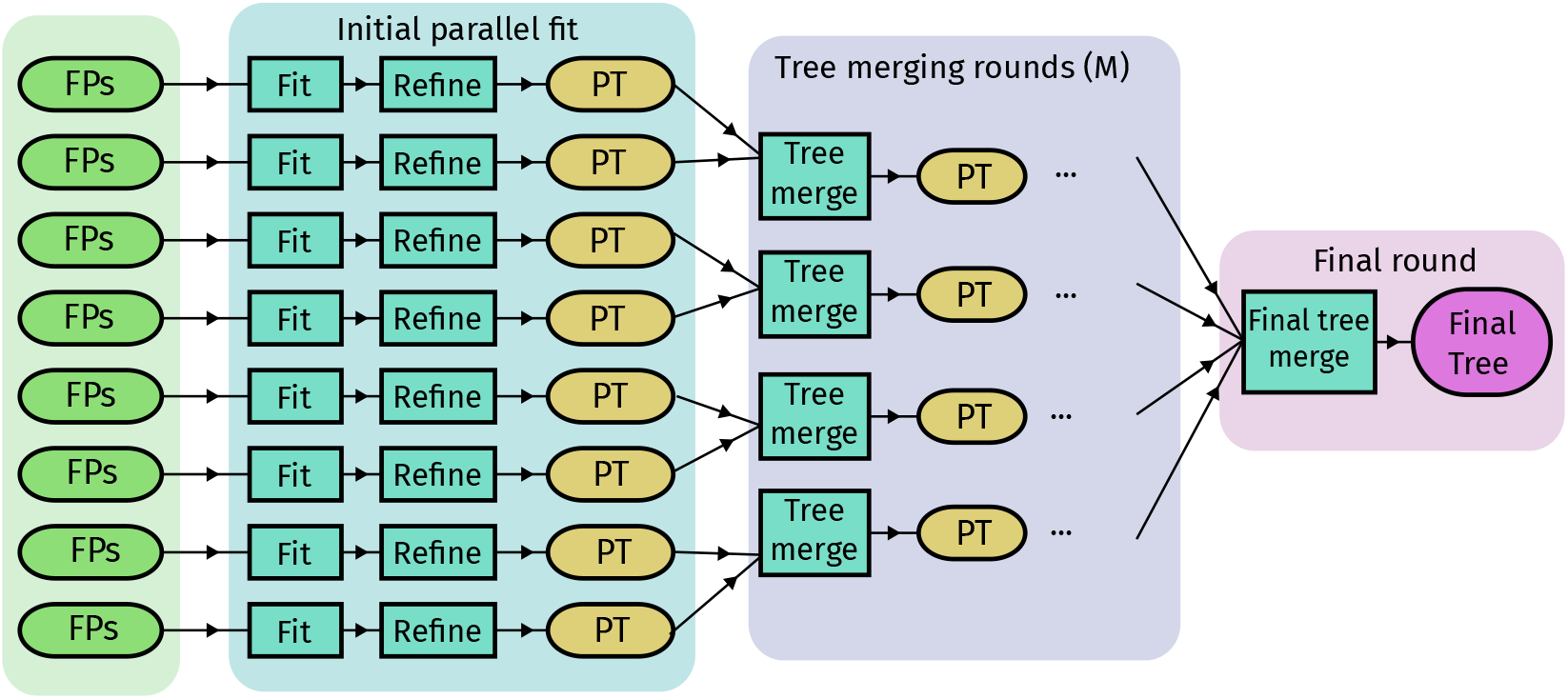
Schematic diagram of the steps in the parallel, multi-round BitBIRCH-Lean algorithm.

In BIRCH-style algorithms the branching factor can significantly affect the performance characteristics, and that is also the case in our parallel, multi-round BitBIRCH implementation. Figure 5 shows the memory usage and total time taken for multi-round BitBIRCH runs when clustering a dataset of 100M molecules, using 30 workers for the initial round and 10 workers for the intermediate tree merging round. In general, we found that a value in the range of 254-1500 for the branching factor is optimal in terms of memory efficiency. This range is robust and only slightly dependent on the features of the database to be clustered. As Figure 5 shows, values on the higher end of this range incur a small performance penalty, but require less RAM usage.

**Figure 5.**
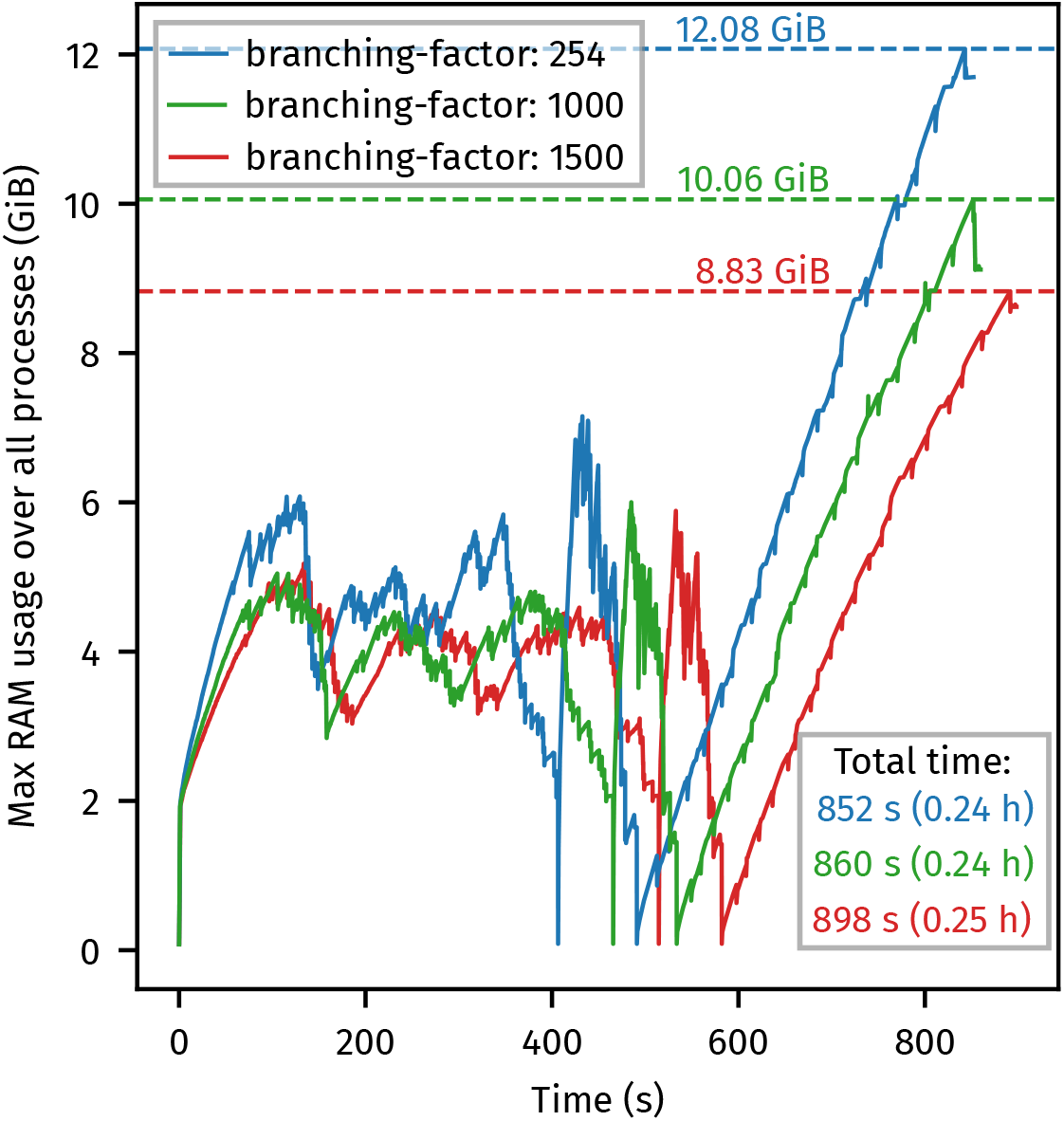
Memory usage and total time for clustering runs of 100M molecules using the parallel, multi-round BitBIRCH-Lean implementation. Runs consist of a single initial run with refinement, a single intermediate tree-merging round, and a final global tree-merging round. All runs use 30 workers for the initial round and 10 workers for the tree-merging round. The *diameter* merge criterion with a threshold value of 0.5 was used in the initial stage of all runs, followed by *tolerance-diameter*, with a tolerance value of 0.05 for tree-merging and refinement.

Figure S4^†^ in the ESI displays the memory usage and total time taken for the same clustering procedure using the serial implementation (branching factor of 1000). The total RAM usage is similar for the serial implementation, since it corresponds to the usage of the final constructed BitBIRCH tree. The total time taken for the serial implementation for the 100M fingerprint set is 2.67 hours, while parallel BitBIRCH clusters the same set in 0.24 hours, 11 times faster.

We investigated the differences between the parallel-multiround and serial BitBIRCH- Lean approaches by clustering a set of 10M fingerprints and comparing the resulting cluster distributions and embeddings through dimensionality reduction. Since in the parallel approach BFs can only merge within a given subsequence, the final clustering is not guaranteed to exactly match the serial results, however, the parallel implementation does allow merging of small clusters in the tree-merging rounds, which reduces the final number of single-element clusters (singletons) and increases the size of clusters with high average Tanimoto similarity (as measured by iSIM).

Figures 6a and 6b show a comparison of the cluster distribution of the top 20 largest clusters for clustering 10M fingerprints using the parallel and serial approaches. Specific values of the top cluster populations and average Tanimoto coefficients can be found in the Table S3^†^ in the ESI. Summary statistics for both distributions are shown in Table S4^†^ in the ESI, which shows that the average cluster size is significantly higher for the parallel case, due to the multiple merging rounds, and that the number of single-element clusters is 3.5 times lower.

**Figure 6.**
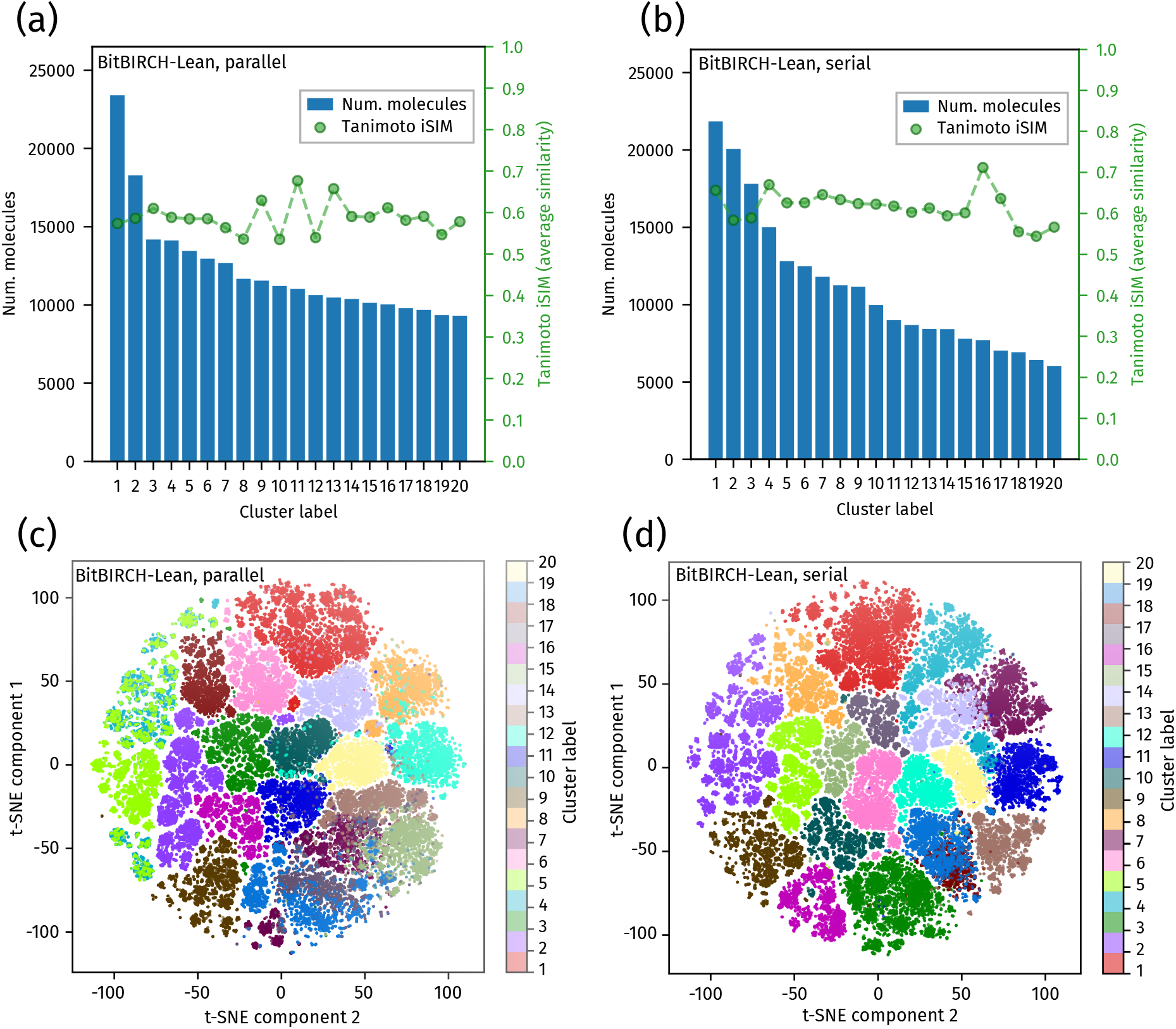
Comparison of clusters populations and within-cluster average Tanimoto similarities (through iSIM values) between parallel (a) and serial (b) BitBIRCH-Lean runs of the sample of 10M fingerprints. PCA-initialized t-SNE visualizations of the top 20 clusters for the same parallel (c) and serial (d) runs are displayed in the bottom panels. A perplexity value of 30 and early-exaggeration of 12 were used in both cases, with a learning rate determined by the dataset size (*N/*12).

The adjusted mutual information score (AMIS)^48^ between two clusterings measures the correspondence of two alternative partitions of the same dataset. Due to the large computational cost of this calculation for a 10M fingerprint set, we estimate the AMIS between the parallel and serial BitBIRCH-Lean partitions by taking the fingerprints corresponding to the 150 most populated clusters of the serial run, resulting in a value of 0.77, indicating significant resemblance in the clustering structure.

Finally, Figures 6c and 6d show a comparison of PCA-initialized t-SNE ^49^ visualizations for both clusterings, which shows that the clustering structure recovered by both schemes is reproduced by the t-SNE clusters. These findings suggest the parallel clustering recovers a very similar structure to the serial approach, with some potential benefits in terms of cluster quality for the parallel case.

### 3.5 Clustering 1 billion molecules in a single workstation

To further demonstrate the scalability and speed of the parallel-multiround BitBIRCH-Lean approach, we apply it to close to 1 billion molecules (954M) from ZINC-22. In order to minimize RAM utilization we use a branching factor of 3500 for this clustering, and perform 4 intermediate *tree-merging* rounds, which helps further coalesce small clusters and singletons. The fingerprint set considered is the same as that used in the original BitBIRCH article. ^30^ Table S5^†^ in the ESI shows the population and average Tanimoto values for this clustering. The largest cluster has a population of around 1.6M molecules, which represents 0.15% of the total number of molecules. This is slightly lower but comparable to the largest cluster for the 100M set, which is in the range of 21000-23000 for the parallel and serial clustering runs, representing 0.22-0.23% of the total. Table S6^†^ shows the corresponding summary statistics. Interestingly, the number of single-element clusters is lower in this case, owing probably to the larger number of tree-merging rounds that enable re-insertion of the singletons into the partial trees.

To cluster this set, the original BitBIRCH parallel approach required more than 2 TiB of RAM and the serial BitBIRCH implementation would have needed multiple days, while relying on indirect label assignment by clustering centroids after an initial parallel round. In contrast, BitBIRCH-Lean’s parallel-multiround approach can cluster it in under 2.5 h, with a memory footprint of 55 GiB of RAM, which is accessible for most modern workstation setups. Figure S4^†^ in the ESI shows the memory usage profile as a function of time for this run. We note that a higher number of processes would increase clustering speed with no total RAM cost, but we opted for 32 processes to better represent the capabilities of typical mid-range workstations.

As a technical note regarding memory usage, we posit that the memory requirements to cluster huge datasets can become less constraining by enabling OS-level memory compression features, for instance, *zswap* ^50,51^ and *zram*.^52,53^ in Linux. The former compresses swap pages in memory, while the latter creates a compressed block device in memory which can be used expand available swap space. Depending on its configuration, *zram* can increase the effective swap capacity tenfold. Other modern operating systems such as Windows and macOS enable analogous features by default, but allow less user control. Preliminary tests with these configurations were promising.

## 4 Conclusions

We introduced *BitBIRCH-Lean*, a memory-efficient, high-throughput implementation of the BitBIRCH clustering algorithm for binary molecular fingerprints. The Lean design centers on two ideas: First, dynamic, safety-checked dtypes for bit feature summaries that avoid premature promotion to 64-bit integers, and second, a packed-bit representation that keeps fingerprints, centroids, and working buffers compact while enabling vectorized bitwise arithmetic. Together, these changes reduce memory usage by more than an order of magnitude relative to dense layouts and preserve the *O*(*N*) scaling critical for very large molecular libraries, and simultaneously increase the algorithm’s performance.

We complemented these changes with improvements in usability and a drop-in *scikitlearn* API, a CLI that exposes full serial and parallel pipelines, and optional C++ kernels (via *pybind11*) for the most expensive inner loops (packed popcounts and centroid similarities). In tests against a state-of-the-art GPU-accelerated implementation of Taylor-Butina, BitBIRCH-Lean achieves substantially lower wall times for large dataset sizes. Peak RAM usage scales linearly, allowing routine processing of hundreds of millions of molecules on a single workstation.

Further, we presented a parallel multi-round strategy that recovers a similar clustering to serial BitBIRCH with a much higher throughput. This implementation greatly improves over the original parallel centroid-clustering approach of BitBIRCH for billion-scale molecular libraries. We observe agreement with serial clustering distributions, and good cluster quality. Future work will explore improved split heuristics, automatic threshold selection, distributed tree-merging schemes, and constant-memory approximations to the parallel multi-round algorithm, which could be optimized using broader quality diagnostics. In terms of performance, more aggressive SIMD specialization and end-to-end GPU implementations may yield further gains.

*BitBIRCH-Lean* is open-source and designed to be practical. By keeping memory footprints small and scaling behavior favorable, it makes clustering of hundred-million-scale libraries accessible on modern workstations, making cheminformatics clustering pipelines widely accessible for researchers even outside traditional HPC setups.

## Supporting information

ESI

## Data and Software Availability

*BitBIRCH-Lean* is open-source software released under the GPL-3.0 license. Code and installation instructions can be found at the GitHub repository https://github.com/mqcomplab/bblean.git. Detailed documentation can be found at https://github.com/mqcomplab/bblean/devdocs.

## Author Contributions

IP: Conceptualization, Methodology, Software, Validation, Formal Analysis, Investigation, Data Curation, Writing, Visualization.

KZ: Software, Formal Analysis, Investigation, Data Curation, Writing.

KLP: Software, Formal Analysis, Investigation, Data Curation, Writing.

ML: Methodology, Software, Validation, Formal Analysis, Investigation, Writing, Visualization.

RAMQ: Conceptualization, Methodology, Software, Validation, Formal Analysis, Investigation, Writing, Supervision, Funding Acquisition.

## Acknowledgement

IP, KZ, KLP, and RAMQ thank and acknowledge funding from the National Institute of General Medical Sciences of the National Institutes of Health, under award R35GM150620. All authors acknowledge technical support from the HiPerGator cluster at the University of Florida.

## Supporting Information Available

Electronic supplementary information (ESI) available.

