## Supplementary material for "BitBIRCH-Lean: chemical space in the palm of your workstation": ESI

#### Contents

**1 Benchmarks for different branchings and C++ extensions S2**

**2 RAM use for serial (100M) and parallel (1B) runs S4**

**3 Detailed benchmarks for BitBIRCH variants S6**

**4 Serial vs. parallel performance: clustering 100M molecules S13**

**5 Statistics and top 20 clusters for clustering 1 billion molecules S14**

### 1 Benchmarks for different branchings and C++ extensions

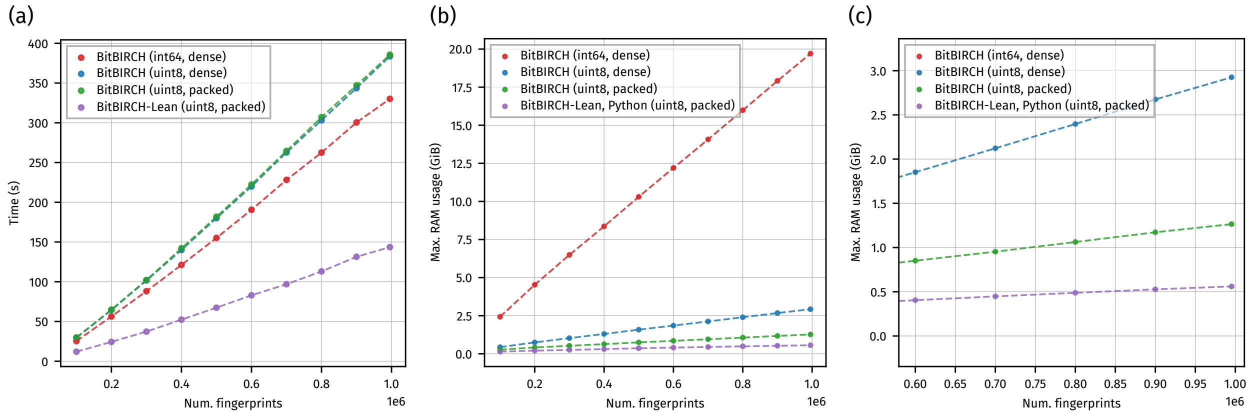


**Figure S1.** Comparison of RAM usage and timings for different BitBIRCH implementations (branching factor 50), for clustering up to 1M molecules. Traces correspond to the average over three independent runs. Traces labeled *int64* correspond to the original BitBIRCH implementation, while traces labeled *uint8* correspond to a naive use of *uint8* arrays without further code modifications. Dense traces use byte-level features in fingerprints, while packed traces correspond to bit-packed fingerprints. All calculations use a threshold of 0.5. (a) presents the timing benchmarks over the full range up to 1M. (b) shows a comparison of the peak RAM usage over this same range, while (c) displays an zoomed-in inset of (b) focusing on the *uint8* and *Lean* implementations, for the largest sets only.

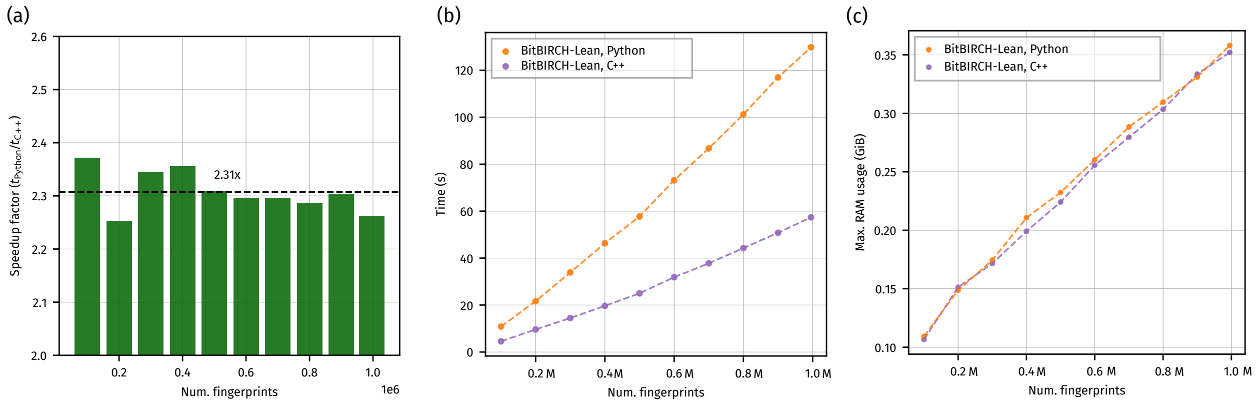


**Figure S2.** Comparison between the Python-only and Python-C++ serial implementations. (a) shows the speedup factor for sets of fingerprints in the range 10k-1M. (b) shows the corresponding timing benchmarks. (c) displays the RAM usage for both implementations (no significant difference between the RAM usage of the Python and Python-C++ implementations was observed).

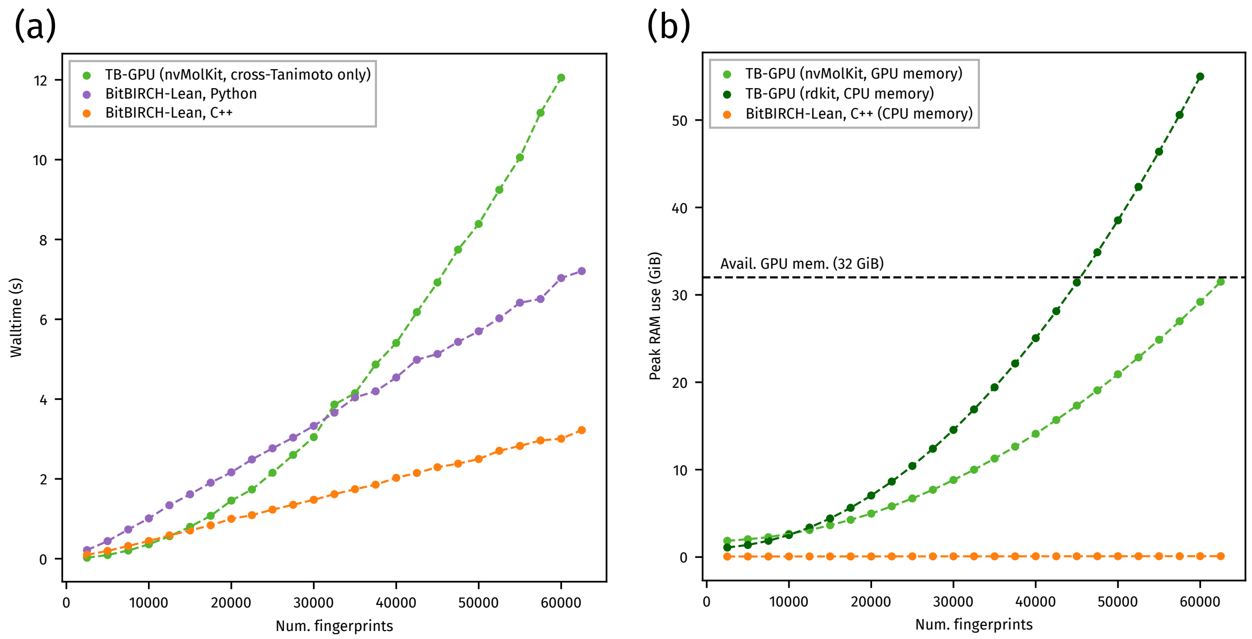


**Figure S3.** Comparison between the GPU-accelerated Taylor-Butina implementation, using nvMolKit with rdkit, and the (serial) BitBIRCH-Lean implementation (both C++ and Python-only). (a) shows timing benchmarks comparing the full BitBIRCH-Lean pipeline with the GPU parallelized cross-Tanimoto kernels of nvMolKit. (b) shows the scaling for memory usage (both CPU and GPU RAM). All benchmarks use a threshold of 0.5.

### 2 RAM use for serial (100M) and parallel (1B) runs

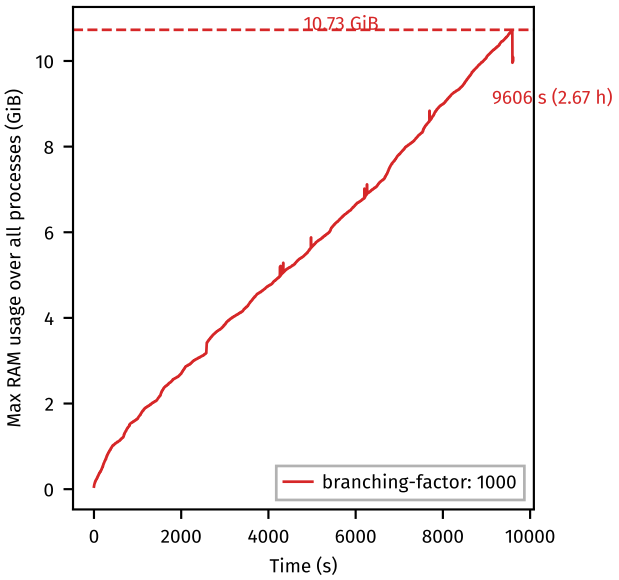


**Figure S4.** Memory usage and total time for clustering a single run (with no refinement) of 100M molecules using the serial BitBIRCH-Lean implementation The *diameter* merge criterion with a threshold value of 0.5, and a branching factor of 1000 were used.

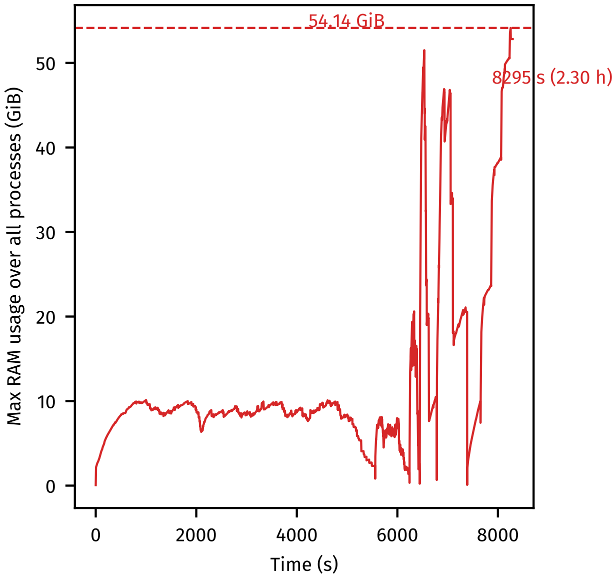


**Figure S5.** Memory usage and total time for clustering a single run of close to 1B molecules (954M), using the parallel, multi-round BitBIRCH-Lean implementation. The run consists of a single initial round with refinement, 4 intermediate tree-merging rounds, and a final global tree-merging round. All runs use 32 workers for the initial round and 12 workers for the tree-merging rounds. The *diameter* merge criterion with a threshold value of 0.5 was used in the initial stage of all runs, followed by *tolerance-diameter*, with a tolerance value of 0.05 for tree-merging and refinement. A branching factor of 3500 was used for this run.

### 3 Detailed benchmarks for BitBIRCH variants

**Table S1.** BitBIRCH-Lean benchmark results using a branching factor of 50

| Variant | Num./1M | Time (s) | Peak RAM use (GiB) |
|---|---|---|---|
| int64-dense | 0.10 | $25.39 \pm 0.24$ | $2.429 \pm 1.91 \times 10^{-3}$ |
| int64-dense | 0.20 | $56.32 \pm 0.04$ | $4.531 \pm 1.35 \times 10^{-3}$ |
| int64-dense | 0.30 | $88.11 \pm 0.50$ | $6.492 \pm 7.21 \times 10^{-4}$ |
| int64-dense | 0.40 | $121.18 \pm 1.11$ | $8.364 \pm 7.66 \times 10^{-4}$ |
| int64-dense | 0.50 | $155.20 \pm 0.55$ | $10.306 \pm 7.06 \times 10^{-5}$ |
| int64-dense | 0.60 | $190.58 \pm 0.35$ | $12.198 \pm 8.53 \times 10^{-4}$ |
| int64-dense | 0.70 | $228.17 \pm 0.73$ | $14.071 \pm 1.31 \times 10^{-3}$ |
| int64-dense | 0.80 | $262.54 \pm 0.41$ | $15.985 \pm 7.02 \times 10^{-4}$ |
| int64-dense | 0.90 | $300.48 \pm 2.06$ | $17.915 \pm 2.74 \times 10^{-4}$ |
| int64-dense | 1.00 | $330.15 \pm 2.18$ | $19.704 \pm 9.49 \times 10^{-4}$ |
| lean-dense | 0.10 | $12.19 \pm 0.18$ | $0.140 \pm 5.10 \times 10^{-4}$ |
| lean-dense | 0.20 | $24.02 \pm 0.11$ | $0.200 \pm 1.48 \times 10^{-3}$ |
| lean-dense | 0.30 | $36.56 \pm 0.04$ | $0.252 \pm 3.61 \times 10^{-3}$ |
| lean-dense | 0.40 | $51.86 \pm 0.48$ | $0.309 \pm 2.94 \times 10^{-4}$ |
| lean-dense | 0.50 | $67.23 \pm 1.40$ | $0.363 \pm 2.65 \times 10^{-3}$ |
| lean-dense | 0.60 | $85.49 \pm 3.61$ | $0.405 \pm 2.44 \times 10^{-4}$ |
| lean-dense | 0.70 | $99.62 \pm 2.99$ | $0.447 \pm 3.06 \times 10^{-4}$ |
| lean-dense | 0.80 | $113.75 \pm 1.85$ | $0.488 \pm 1.55 \times 10^{-4}$ |
| lean-dense | 0.90 | $128.54 \pm 1.78$ | $0.527 \pm 6.42 \times 10^{-4}$ |
| lean-dense | 1.00 | $144.77 \pm 3.44$ | $0.560 \pm 4.44 \times 10^{-4}$ |
| lean-dense-cpp | 0.10 | $5.49 \pm 0.07$ | $0.141 \pm 2.74 \times 10^{-3}$ |

Continued on next page

**Table S1.** BitBIRCH-Lean benchmark results using a branching factor of 50

| Variant | Num./1M | Time (s) | Peak RAM use (GiB) |
|---|---|---|---|
| lean-dense-cpp | 0.20 | $11.21 \pm 0.11$ | $0.206 \pm 4.70 \times 10^{-3}$ |
| lean-dense-cpp | 0.30 | $17.12 \pm 0.09$ | $0.253 \pm 2.98 \times 10^{-3}$ |
| lean-dense-cpp | 0.40 | $23.94 \pm 0.45$ | $0.307 \pm 9.39 \times 10^{-4}$ |
| lean-dense-cpp | 0.50 | $30.90 \pm 0.30$ | $0.364 \pm 6.26 \times 10^{-4}$ |
| lean-dense-cpp | 0.60 | $37.86 \pm 0.36$ | $0.404 \pm 9.11 \times 10^{-4}$ |
| lean-dense-cpp | 0.70 | $45.12 \pm 0.36$ | $0.446 \pm 1.44 \times 10^{-3}$ |
| lean-dense-cpp | 0.80 | $54.54 \pm 2.16$ | $0.489 \pm 2.09 \times 10^{-4}$ |
| lean-dense-cpp | 0.90 | $60.69 \pm 1.15$ | $0.527 \pm 7.55 \times 10^{-4}$ |
| lean-dense-cpp | 1.00 | $68.26 \pm 2.05$ | $0.558 \pm 1.19 \times 10^{-4}$ |
| lean-packed | 0.10 | $11.99 \pm 0.05$ | $0.140 \pm 2.83 \times 10^{-4}$ |
| lean-packed | 0.20 | $24.33 \pm 0.42$ | $0.207 \pm 5.30 \times 10^{-3}$ |
| lean-packed | 0.30 | $37.25 \pm 0.58$ | $0.254 \pm 1.92 \times 10^{-3}$ |
| lean-packed | 0.40 | $52.37 \pm 0.44$ | $0.307 \pm 6.60 \times 10^{-4}$ |
| lean-packed | 0.50 | $67.34 \pm 0.70$ | $0.363 \pm 6.96 \times 10^{-4}$ |
| lean-packed | 0.60 | $82.97 \pm 0.71$ | $0.405 \pm 8.22 \times 10^{-4}$ |
| lean-packed | 0.70 | $96.90 \pm 0.53$ | $0.447 \pm 3.35 \times 10^{-4}$ |
| lean-packed | 0.80 | $113.01 \pm 0.50$ | $0.488 \pm 4.44 \times 10^{-4}$ |
| lean-packed | 0.90 | $131.38 \pm 3.15$ | $0.528 \pm 4.58 \times 10^{-4}$ |
| lean-packed | 1.00 | $143.58 \pm 2.51$ | $0.560 \pm 5.31 \times 10^{-4}$ |
| lean-packed-cpp | 0.10 | $5.42 \pm 0.20$ | $0.142 \pm 6.16 \times 10^{-4}$ |
| lean-packed-cpp | 0.20 | $10.85 \pm 0.12$ | $0.201 \pm 7.00 \times 10^{-3}$ |
| lean-packed-cpp | 0.30 | $16.43 \pm 0.05$ | $0.248 \pm 2.27 \times 10^{-3}$ |
| lean-packed-cpp | 0.40 | $23.27 \pm 0.44$ | $0.300 \pm 2.38 \times 10^{-3}$ |

Continued on next page

**Table S1.** BitBIRCH-Lean benchmark results using a branching factor of 50

| Variant | Num./1M | Time (s) | Peak RAM use (GiB) |
|---|---|---|---|
| lean-packed-cpp | 0.50 | $29.91 \pm 0.41$ | $0.355 \pm 3.81 \times 10^{-3}$ |
| lean-packed-cpp | 0.60 | $36.13 \pm 0.37$ | $0.394 \pm 1.99 \times 10^{-3}$ |
| lean-packed-cpp | 0.70 | $44.21 \pm 1.12$ | $0.437 \pm 1.58 \times 10^{-4}$ |
| lean-packed-cpp | 0.80 | $50.38 \pm 1.11$ | $0.477 \pm 6.88 \times 10^{-4}$ |
| lean-packed-cpp | 0.90 | $57.45 \pm 0.26$ | $0.515 \pm 9.28 \times 10^{-4}$ |
| lean-packed-cpp | 1.00 | $63.57 \pm 0.54$ | $0.547 \pm 7.18 \times 10^{-4}$ |
| uint8-dense | 0.10 | $29.68 \pm 0.28$ | $0.437 \pm 7.99 \times 10^{-4}$ |
| uint8-dense | 0.20 | $64.42 \pm 0.63$ | $0.747 \pm 2.42 \times 10^{-4}$ |
| uint8-dense | 0.30 | $101.70 \pm 0.50$ | $1.028 \pm 3.30 \times 10^{-4}$ |
| uint8-dense | 0.40 | $140.00 \pm 0.23$ | $1.301 \pm 3.59 \times 10^{-4}$ |
| uint8-dense | 0.50 | $180.09 \pm 0.42$ | $1.583 \pm 6.88 \times 10^{-4}$ |
| uint8-dense | 0.60 | $219.83 \pm 0.31$ | $1.852 \pm 6.69 \times 10^{-4}$ |
| uint8-dense | 0.70 | $262.71 \pm 0.57$ | $2.122 \pm 5.28 \times 10^{-4}$ |
| uint8-dense | 0.80 | $303.25 \pm 0.97$ | $2.397 \pm 5.06 \times 10^{-4}$ |
| uint8-dense | 0.90 | $343.61 \pm 2.30$ | $2.675 \pm 7.89 \times 10^{-4}$ |
| uint8-dense | 1.00 | $383.68 \pm 0.49$ | $2.925 \pm 4.46 \times 10^{-4}$ |
| uint8-packed | 0.10 | $29.74 \pm 0.05$ | $0.270 \pm 3.42 \times 10^{-4}$ |
| uint8-packed | 0.20 | $64.87 \pm 0.36$ | $0.413 \pm 3.56 \times 10^{-4}$ |
| uint8-packed | 0.30 | $102.08 \pm 0.25$ | $0.527 \pm 4.95 \times 10^{-4}$ |
| uint8-packed | 0.40 | $142.03 \pm 0.40$ | $0.634 \pm 4.18 \times 10^{-4}$ |
| uint8-packed | 0.50 | $181.77 \pm 0.89$ | $0.749 \pm 2.23 \times 10^{-4}$ |
| uint8-packed | 0.60 | $222.38 \pm 0.34$ | $0.851 \pm 8.37 \times 10^{-5}$ |
| uint8-packed | 0.70 | $264.56 \pm 1.24$ | $0.953 \pm 9.92 \times 10^{-5}$ |

Continued on next page

**Table S1.** BitBIRCH-Lean benchmark results using a branching factor of 50

| Variant | Num./1M | Time (s) | Peak RAM use (GiB) |
|---|---|---|---|
| uint8-packed | 0.80 | $307.08 \pm 0.91$ | $1.063 \pm 4.92 \times 10^{-4}$ |
| uint8-packed | 0.90 | $347.10 \pm 3.68$ | $1.173 \pm 4.56 \times 10^{-4}$ |
| uint8-packed | 1.00 | $385.68 \pm 1.32$ | $1.265 \pm 2.61 \times 10^{-4}$ |

**Table S2.** BitBIRCH-Lean benchmark results using a branching factor of 254

| Variant | Num./1M | Time (s) | Peak RAM use (GiB) |
|---|---|---|---|
| int64-dense | 0.10 | $57.44 \pm 0.39$ | $2.124 \pm 6.26 \times 10^{-4}$ |
| int64-dense | 0.20 | $133.70 \pm 0.06$ | $3.992 \pm 8.65 \times 10^{-4}$ |
| int64-dense | 0.30 | $220.32 \pm 0.35$ | $5.769 \pm 5.97 \times 10^{-4}$ |
| int64-dense | 0.40 | $310.41 \pm 0.60$ | $7.563 \pm 2.91 \times 10^{-4}$ |
| int64-dense | 0.50 | $412.35 \pm 1.06$ | $9.296 \pm 4.26 \times 10^{-4}$ |
| int64-dense | 0.60 | $497.51 \pm 0.57$ | $11.054 \pm 4.89 \times 10^{-4}$ |
| int64-dense | 0.70 | $589.47 \pm 0.74$ | $12.798 \pm 1.62 \times 10^{-4}$ |
| int64-dense | 0.80 | $680.55 \pm 0.90$ | $14.516 \pm 4.25 \times 10^{-4}$ |
| int64-dense | 0.90 | $775.74 \pm 2.99$ | $16.213 \pm 5.23 \times 10^{-4}$ |
| int64-dense | 1.00 | $867.79 \pm 0.95$ | $17.791 \pm 3.02 \times 10^{-4}$ |
| lean-dense | 0.10 | $10.50 \pm 0.12$ | $0.112 \pm 3.18 \times 10^{-3}$ |
| lean-dense | 0.20 | $21.70 \pm 0.53$ | $0.149 \pm 3.15 \times 10^{-4}$ |
| lean-dense | 0.30 | $33.18 \pm 0.14$ | $0.175 \pm 5.22 \times 10^{-3}$ |
| lean-dense | 0.40 | $45.63 \pm 0.42$ | $0.204 \pm 1.64 \times 10^{-3}$ |
| lean-dense | 0.50 | $58.21 \pm 1.39$ | $0.236 \pm 1.57 \times 10^{-3}$ |
| lean-dense | 0.60 | $72.65 \pm 1.03$ | $0.261 \pm 1.24 \times 10^{-3}$ |

Continued on next page

**Table S2.** BitBIRCH-Lean benchmark results using a branching factor of 254

| Variant | Num./1M | Time (s) | Peak RAM use (GiB) |
|---|---|---|---|
| lean-dense | 0.70 | $86.02 \pm 0.31$ | $0.288 \pm 1.33 \times 10^{-3}$ |
| lean-dense | 0.80 | $99.79 \pm 0.93$ | $0.310 \pm 3.42 \times 10^{-4}$ |
| lean-dense | 0.90 | $114.42 \pm 0.55$ | $0.338 \pm 4.47 \times 10^{-4}$ |
| lean-dense | 1.00 | $129.90 \pm 2.20$ | $0.356 \pm 5.12 \times 10^{-4}$ |
| lean-dense-cpp | 0.10 | $4.86 \pm 0.09$ | $0.110 \pm 3.33 \times 10^{-3}$ |
| lean-dense-cpp | 0.20 | $9.77 \pm 0.04$ | $0.152 \pm 4.03 \times 10^{-3}$ |
| lean-dense-cpp | 0.30 | $15.12 \pm 0.15$ | $0.180 \pm 2.94 \times 10^{-3}$ |
| lean-dense-cpp | 0.40 | $20.39 \pm 0.38$ | $0.211 \pm 5.22 \times 10^{-3}$ |
| lean-dense-cpp | 0.50 | $25.68 \pm 0.07$ | $0.230 \pm 6.58 \times 10^{-3}$ |
| lean-dense-cpp | 0.60 | $32.54 \pm 0.61$ | $0.261 \pm 3.99 \times 10^{-3}$ |
| lean-dense-cpp | 0.70 | $39.38 \pm 0.48$ | $0.287 \pm 2.49 \times 10^{-3}$ |
| lean-dense-cpp | 0.80 | $45.61 \pm 0.52$ | $0.310 \pm 1.06 \times 10^{-3}$ |
| lean-dense-cpp | 0.90 | $52.17 \pm 0.17$ | $0.332 \pm 5.07 \times 10^{-3}$ |
| lean-dense-cpp | 1.00 | $59.27 \pm 0.74$ | $0.355 \pm 1.53 \times 10^{-3}$ |
| lean-packed | 0.10 | $10.85 \pm 0.42$ | $0.111 \pm 1.87 \times 10^{-3}$ |
| lean-packed | 0.20 | $21.66 \pm 0.10$ | $0.151 \pm 3.90 \times 10^{-3}$ |
| lean-packed | 0.30 | $33.90 \pm 0.38$ | $0.174 \pm 4.91 \times 10^{-3}$ |
| lean-packed | 0.40 | $46.33 \pm 1.40$ | $0.211 \pm 6.41 \times 10^{-3}$ |
| lean-packed | 0.50 | $57.71 \pm 0.61$ | $0.236 \pm 1.99 \times 10^{-3}$ |
| lean-packed | 0.60 | $73.14 \pm 1.27$ | $0.266 \pm 1.49 \times 10^{-4}$ |
| lean-packed | 0.70 | $86.73 \pm 1.57$ | $0.288 \pm 1.68 \times 10^{-3}$ |
| lean-packed | 0.80 | $101.20 \pm 1.78$ | $0.310 \pm 1.61 \times 10^{-3}$ |
| lean-packed | 0.90 | $116.95 \pm 2.36$ | $0.331 \pm 4.81 \times 10^{-3}$ |

Continued on next page

**Table S2.** BitBIRCH-Lean benchmark results using a branching factor of 254

| Variant | Num./1M | Time (s) | Peak RAM use (GiB) |
|---|---|---|---|
| lean-packed | 1.00 | $129.80 \pm 0.80$ | $0.358 \pm 3.95 \times 10^{-3}$ |
| lean-packed-cpp | 0.10 | $4.57 \pm 0.03$ | $0.110 \pm 2.47 \times 10^{-3}$ |
| lean-packed-cpp | 0.20 | $9.62 \pm 0.30$ | $0.152 \pm 6.51 \times 10^{-4}$ |
| lean-packed-cpp | 0.30 | $14.46 \pm 0.25$ | $0.175 \pm 5.82 \times 10^{-3}$ |
| lean-packed-cpp | 0.40 | $19.66 \pm 0.08$ | $0.204 \pm 6.97 \times 10^{-3}$ |
| lean-packed-cpp | 0.50 | $25.00 \pm 0.32$ | $0.232 \pm 1.77 \times 10^{-3}$ |
| lean-packed-cpp | 0.60 | $31.86 \pm 0.30$ | $0.256 \pm 8.83 \times 10^{-4}$ |
| lean-packed-cpp | 0.70 | $37.78 \pm 0.20$ | $0.282 \pm 1.77 \times 10^{-3}$ |
| lean-packed-cpp | 0.80 | $44.27 \pm 0.56$ | $0.303 \pm 1.44 \times 10^{-3}$ |
| lean-packed-cpp | 0.90 | $50.79 \pm 0.31$ | $0.334 \pm 8.97 \times 10^{-3}$ |
| lean-packed-cpp | 1.00 | $57.37 \pm 1.05$ | $0.352 \pm 2.35 \times 10^{-3}$ |
| uint8-dense | 0.10 | $58.34 \pm 0.04$ | $0.378 \pm 4.47 \times 10^{-4}$ |
| uint8-dense | 0.20 | $132.65 \pm 0.48$ | $0.637 \pm 2.74 \times 10^{-4}$ |
| uint8-dense | 0.30 | $216.89 \pm 1.13$ | $0.881 \pm 1.33 \times 10^{-4}$ |
| uint8-dense | 0.40 | $308.87 \pm 1.28$ | $1.127 \pm 5.30 \times 10^{-4}$ |
| uint8-dense | 0.50 | $411.41 \pm 0.34$ | $1.361 \pm 7.84 \times 10^{-4}$ |
| uint8-dense | 0.60 | $495.90 \pm 0.49$ | $1.606 \pm 4.37 \times 10^{-4}$ |
| uint8-dense | 0.70 | $580.45 \pm 1.46$ | $1.842 \pm 2.85 \times 10^{-4}$ |
| uint8-dense | 0.80 | $670.49 \pm 2.26$ | $2.072 \pm 8.78 \times 10^{-5}$ |
| uint8-dense | 0.90 | $764.39 \pm 1.61$ | $2.300 \pm 6.41 \times 10^{-4}$ |
| uint8-dense | 1.00 | $854.69 \pm 2.21$ | $2.505 \pm 3.41 \times 10^{-4}$ |
| uint8-packed | 0.10 | $58.19 \pm 0.21$ | $0.212 \pm 2.00 \times 10^{-4}$ |
| uint8-packed | 0.20 | $132.81 \pm 0.12$ | $0.304 \pm 7.45 \times 10^{-4}$ |

Continued on next page

**Table S2.** BitBIRCH-Lean benchmark results using a branching factor of 254

| Variant | Num./1M | Time (s) | Peak RAM use (GiB) |
|---|---|---|---|
| uint8-packed | 0.30 | $217.78 \pm 0.26$ | $0.380 \pm 3.16 \times 10^{-4}$ |
| uint8-packed | 0.40 | $311.02 \pm 1.97$ | $0.460 \pm 5.11 \times 10^{-4}$ |
| uint8-packed | 0.50 | $412.00 \pm 1.12$ | $0.527 \pm 4.81 \times 10^{-4}$ |
| uint8-packed | 0.60 | $497.97 \pm 1.34$ | $0.605 \pm 2.54 \times 10^{-4}$ |
| uint8-packed | 0.70 | $582.47 \pm 0.83$ | $0.674 \pm 7.54 \times 10^{-4}$ |
| uint8-packed | 0.80 | $672.50 \pm 1.51$ | $0.737 \pm 6.86 \times 10^{-4}$ |
| uint8-packed | 0.90 | $767.51 \pm 1.82$ | $0.798 \pm 7.06 \times 10^{-5}$ |
| uint8-packed | 1.00 | $857.73 \pm 1.82$ | $0.845 \pm 3.56 \times 10^{-4}$ |

### 4 Serial vs. parallel performance: clustering 100M molecules

**Table S3.** Top 20 clusters: comparison between Serial and Parallel.

| Serial | | | Parallel | | |
|---|---|---|---|---|---|
| Cluster size | % Fingerprints | iSIM | Cluster size | % Fingerprints | iSIM |
| 21848 | 0.22 | 0.656 | 23404 | 0.23 | 0.574 |
| 20072 | 0.20 | 0.583 | 18277 | 0.18 | 0.586 |
| 17809 | 0.18 | 0.589 | 14182 | 0.14 | 0.610 |
| 15002 | 0.15 | 0.670 | 14118 | 0.14 | 0.589 |
| 12816 | 0.13 | 0.626 | 13447 | 0.13 | 0.585 |
| 12489 | 0.13 | 0.626 | 12957 | 0.13 | 0.585 |
| 11799 | 0.12 | 0.646 | 12672 | 0.13 | 0.564 |
| 11255 | 0.11 | 0.634 | 11669 | 0.12 | 0.536 |
| 11163 | 0.11 | 0.624 | 11551 | 0.12 | 0.630 |
| 9972 | 0.10 | 0.622 | 11214 | 0.11 | 0.535 |
| 8997 | 0.09 | 0.618 | 11022 | 0.11 | 0.677 |
| 8680 | 0.09 | 0.603 | 10631 | 0.11 | 0.540 |
| 8425 | 0.08 | 0.613 | 10470 | 0.10 | 0.658 |
| 8412 | 0.08 | 0.594 | 10381 | 0.10 | 0.591 |
| 7795 | 0.08 | 0.601 | 10132 | 0.10 | 0.589 |
| 7708 | 0.08 | 0.712 | 10031 | 0.10 | 0.612 |
| 7031 | 0.07 | 0.636 | 9788 | 0.10 | 0.582 |
| 6924 | 0.07 | 0.555 | 9680 | 0.10 | 0.591 |
| 6419 | 0.06 | 0.544 | 9342 | 0.09 | 0.547 |
| 6035 | 0.06 | 0.566 | 9309 | 0.09 | 0.578 |

**Table S4.** Summary statistics: comparison between Serial and Parallel.

| Summary metric | Serial | Parallel |
|---|---|---|
| Total num. fps | 9,977,141 | 9,977,141 |
| Total num. clusters | 420,622 | 243,615 |
| Total num. singletons | 123,126 (29.27%) | 34,745 (14.26%) |
| Clusters with size > 10 | 117,166 (27.86%) | 89,550 (36.76%) |
| Clusters with size > 100 | 19,186 (4.56%) | 19,486 (8.00%) |
| Num-clusters / num-fps ratio | 0.04 | 0.02 |
| Mean size | 23.72 | 40.95 |
| Max. size | 21,848 | 23,404 |

### 5 Statistics and top 20 clusters for clustering 1 billion molecules

**Table S5.** Top 20 clusters: 1 billion molecules from ZINC-22

| Cluster size | % Fingerprints | iSIM |
|---|---|---|
| 1586115 | 0.17 | 0.523 |
| 1144078 | 0.12 | 0.546 |
| 969766 | 0.10 | 0.633 |
| 630870 | 0.07 | 0.568 |
| 608528 | 0.06 | 0.655 |
| 600713 | 0.06 | 0.549 |
| 524324 | 0.05 | 0.604 |
| 503723 | 0.05 | 0.526 |
| 464688 | 0.05 | 0.562 |
| 445101 | 0.05 | 0.542 |
| 428567 | 0.04 | 0.589 |
| 399770 | 0.04 | 0.627 |
| 393878 | 0.04 | 0.570 |
| 388242 | 0.04 | 0.548 |
| 374118 | 0.04 | 0.531 |
| 355374 | 0.04 | 0.639 |
| 352404 | 0.04 | 0.525 |
| 324158 | 0.03 | 0.580 |
| 321064 | 0.03 | 0.625 |
| 319544 | 0.03 | 0.695 |

**Table S6.** Summary statistics: Clustering 1 billion molecules from ZINC-22

| Summary metric | Value |
|---|---|
| Total num. fps | 954,748,201 |
| Total num. clusters | 2,654,797 |
| Total num. singletons | 55,223 (2.08%) |
| Clusters with size $>$ 10 | 1,785,392 (67.25%) |
| Clusters with size $>$ 100 | 834,471 (31.43%) |
| Num-clusters / num-fps ratio | 0.00 |
| Mean size | 359.63 |
| Max. size | 1,586,115 |
